# NeoCoMM: Neocortical Computational Microscale Model

**DOI:** 10.1101/2024.04.08.588273

**Authors:** M. Al Harrach, M. Yochum, F. Wendling

## Abstract

The Neocortical Computational Microscale model (NeoCoMM) is a unique neurophysiologically-inspired software. It offers a friendly graphical user interface that allows for the simulation of the intracellular and extracellular neural activity of a neocortical column. This software provides a realistic framework that can portray the neural activity and underlying cellular mechanisms related to different brain pathologies such as epilepsy. NeoCoMM is capable of (1) simulating the cortical tissue of three different species, (2) visualizing individual cell responses to external stimulation, (3) visualizing the corresponding local field potential, (4) studying the impact of the recording electrode features on simulated signals, and (5) testing various physiological and pathological hypotheses. While NeoCoMM was primarily developed for simulating epileptiform activity, it can also generate healthy brain rhythms or be adapted to other brain disorders.

## 1. Motivation and significance

Computational models of the brain are opening new avenues for experimental and clinical research [1]. These models are capable of depicting the dynamics of the brain across a spectrum of scales, encompassing cellular (microscopic), population (neural mass), and whole-brain (macroscopic) levels [2]. In particular, microscale models can be a valuable tool for understanding underlying physiological and pathological mechanisms [3, 4, 5, 6]. They offer insights into the cellular dynamics of neural networks in various contexts, especially in the case of brain disorders such as epilepsy [5, 7, 8]. These models need to be neurophysiologically and biophysically relevant which can be achieved thanks to recent advances in computation and increased experimental data gathering [4]. An example of these models would be the Blue Brain Project’s study where Markram et al. presented a new detailed reconstruction and simulation of the neocortical microcircuitry [9]. As another example, Schneider et al. introduced a full-scale computational model of the dentate gyrus that is highly detailed [10]. However, despite their promising detailed approaches and highly realistic representation of the brain circuits, these kinds of models require high-performance computing resources to be used not to mention the need for compatible hardware to match the complexity of the algorithm kernels [4]. Moreover, the creation of large networks from these models still requires a series of approximations [6].

**Table 1:** Code metadata.

| Nr. | Code metadata description | Please fill in this column |
| --- | --- | --- |
| C1 | Current code version | V1 |
| C2 | Permanent link to code/repository used for this code version | For example: https://github.com/MariamAlHarrach/NeoCoMM |
| C3 | Permanent link to Reproducible Capsule |  |
| C4 | Legal Code License | MIT License |
| C5 | Code versioning system used | git |
| C6 | Software code languages, tools, and services used | python3.10 or 3.11 |
| C7 | Compilation requirements, operating environments & dependencies | matplotlib3.8.2, scipy1.12.0, numpy1.26.3 numba0.58.1, vtk9.3.0, pyqt55.15.10,pyedflib0.1.36 |
| C8 | If available Link to developer documentation/manual | https://github.com/MariamAlHarrach/NeoCoMM/blob/main/NeoCoMM_Manual_V0.0.docx |
| C9 | Support email for questions |  |

In this paper, we present a novel microscale simplified model of cortical circuitry that draws inspiration from neuroscience and is both physiologically and biophysically credible. It allows for the simulation of neural network activity at the intracellular and extracellular levels (Local field potentials (LFPs)). The model’s structural (multilayered structure, geometrical features, cell distribution, connectivity) and electrophysiological characteristics were based on real data (experimental recordings). The presented software can be used as a tool for hypothesis testing and exploration. In addition, it offers a framework for developing new models with multiple cortical networks (columns). The model also permits the simulation of the cortical column for three different species: Humans, rats, and mice. Another novelty that is included in the software is the electrode-tissue interface that allows the user to simulate realistic LFPs recorded by a platinum SEEG electrode in the case of human recordings and a wire-type stainless steel electrode in the case of rat and mouse simulations. The electrode shape, position, and orientation can also be adjusted and tested by the user.

The NeoCoMM software can be an asset for researchers investigating different brain disorders. In the case of epilepsy, NeoCoMM has been recently used to create an epileptic neocortical column by adjusting the electrophysiological parameters of different cell types (synaptic receptors conductance values) to render the tissue hyperexcitable [11]. The new epileptic configuration was then used to simulate different types of epileptic events (interictal spikes, spike waves, polyspike waves, and high-frequency oscillations). Taking advantage of the simultaneous representation of intracellular and extracellular activity provided by NeoCOMM, we were able to uncover the underlying cellular dynamics responsible for each type of interictal event along with the corresponding parameters responsible for these dynamics creating the epileptic event. This was done for both human and mouse simulations and was compared to real clinical and experimental signals respectively. Finally, the study also investigated the impact of the electrode’s geometrical characteristics on the simulated LFP [11].

## 2. Software description

### 2.1. Software architecture

The software was designed to be used with a graphical user interface (Figure 1), which allows users to operate the model without the need for programming. Additional panels are also included in the software giving the user full control over the cortical tissue configuration and modification (Figure 2). Nevertheless, the software remains accessible to expert programmers who can choose to employ scripts for their purposes making easier the use of automated routines. An example of such a script is included in the software package (file Script_automate.py at https://gitlab.univ-rennes1.fr/myochum/neocomm).

**Figure 1:**
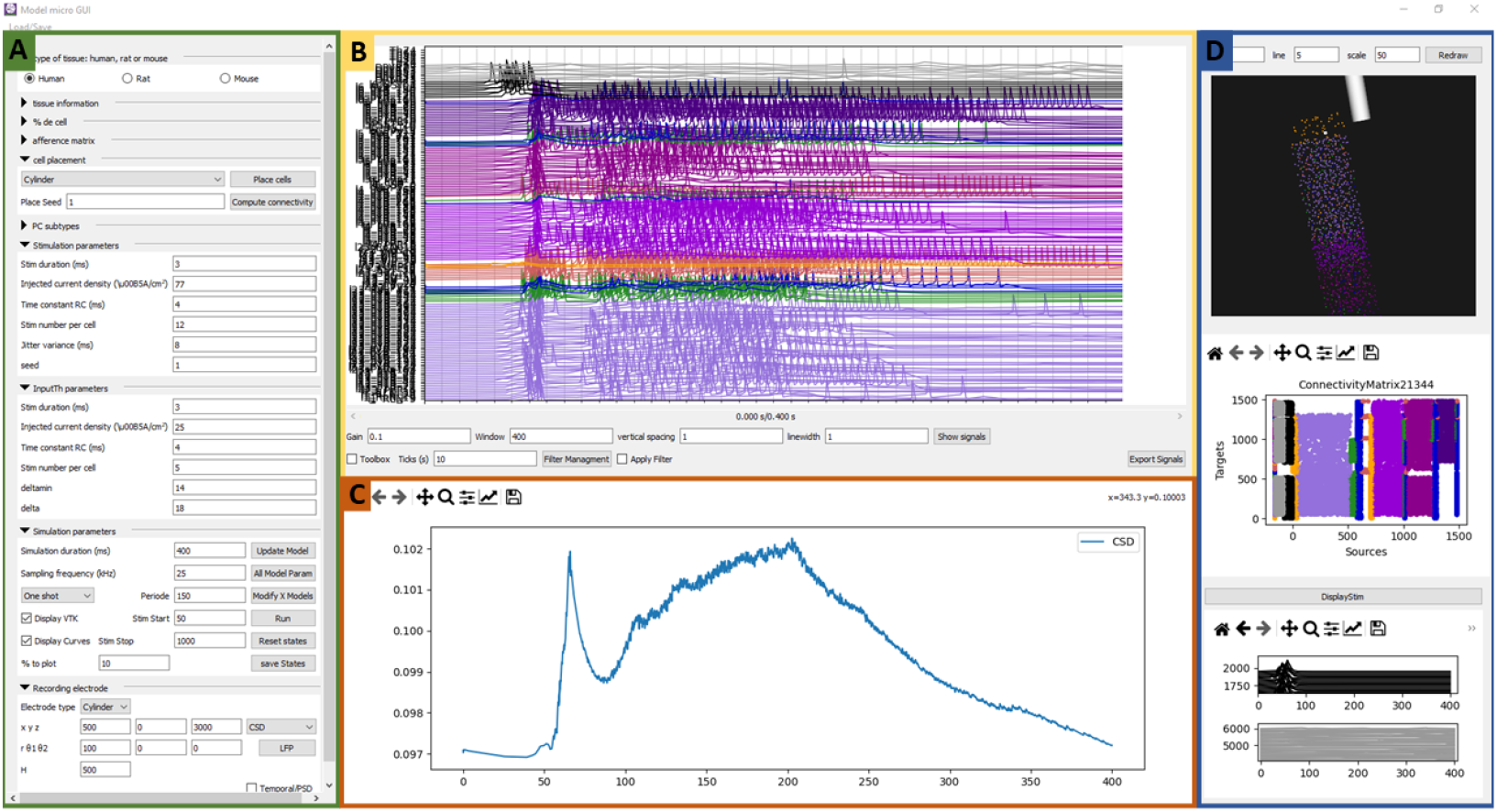
The graphical user interface of NeoCoMM. **A:** The cortical column parameter configuration panel. This panel includes The type of tissue, tissue information, cell distribution percentage, connectivity afference matrix, cell placement type, PC subtypes distribution, external input stimulation parameters, simulation parameters, and the recording electrode characteristics parameters. **B:** The intracellular activity representation panel. **C:** The local field potential plot. **D:** The 3D cortical tissue representation with the corresponding connectivity matrix as well as the stimulation signals plot panel.

**Figure 2:**
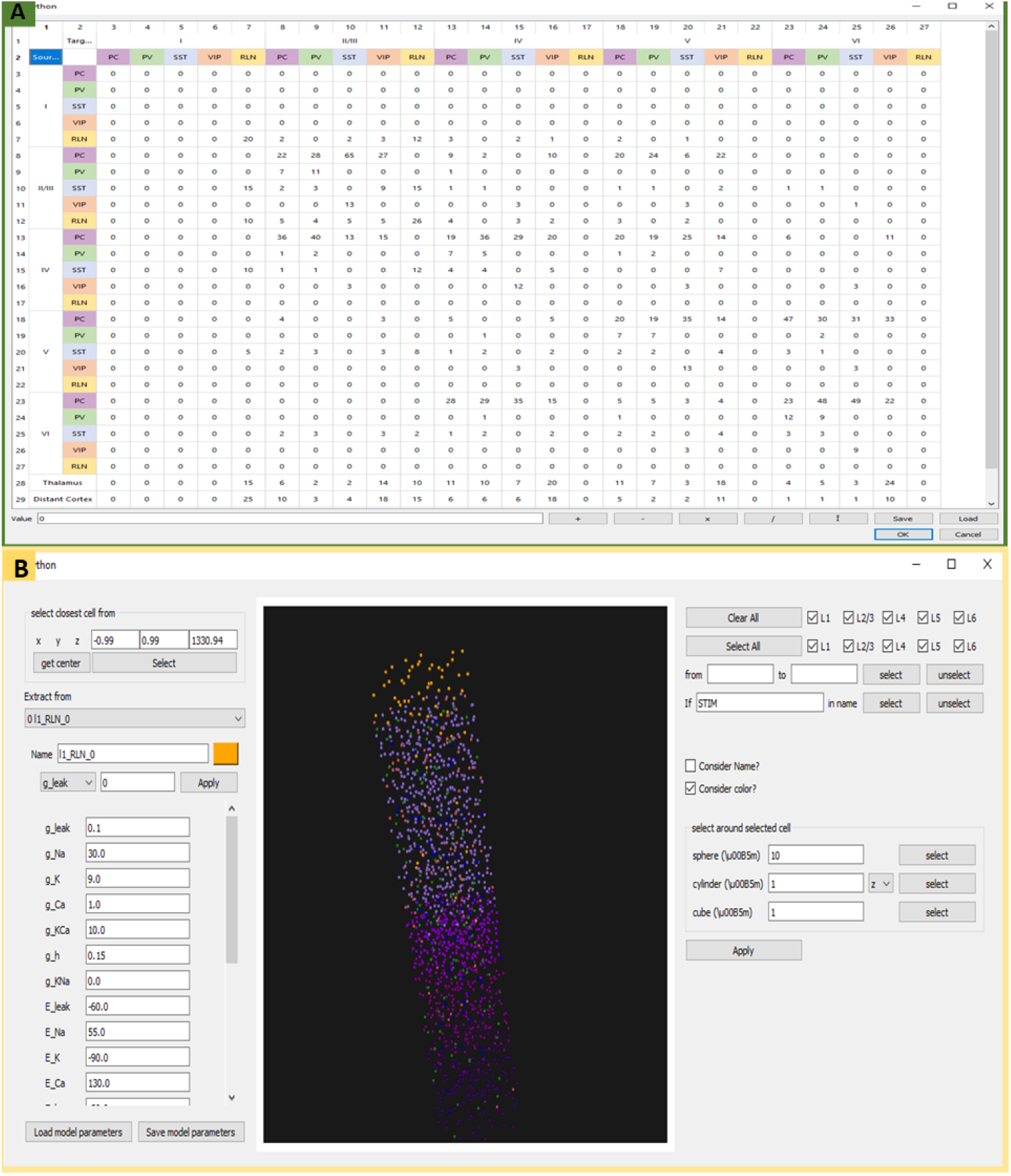
Additional panels of the NeoCoMM graphical user interface. **A:** The afference matrix panel. Each column presents the percentage of afference connections per cell type. It allows the loading of an already existing matrix or the modification of the default matrix values. **B:** The “modify tissue parameter” panel. This panel allows the user to modify the physiological parameters of the neural network. The user can define a specific cluster of a predefined shape (sphere, cylinder, or cube) and select a specific cell type to adjust its parameters. The user can also select cells based on their types or their position in the layers, their name, or their number. The modifications can either be applied to the parameters of one selected cell or to a cluster of selected cells.

The architecture of cortical tissue creation and configuration is shown in Figure 3. It depicts the implementation sequence that comprises of:

**Figure 3:**
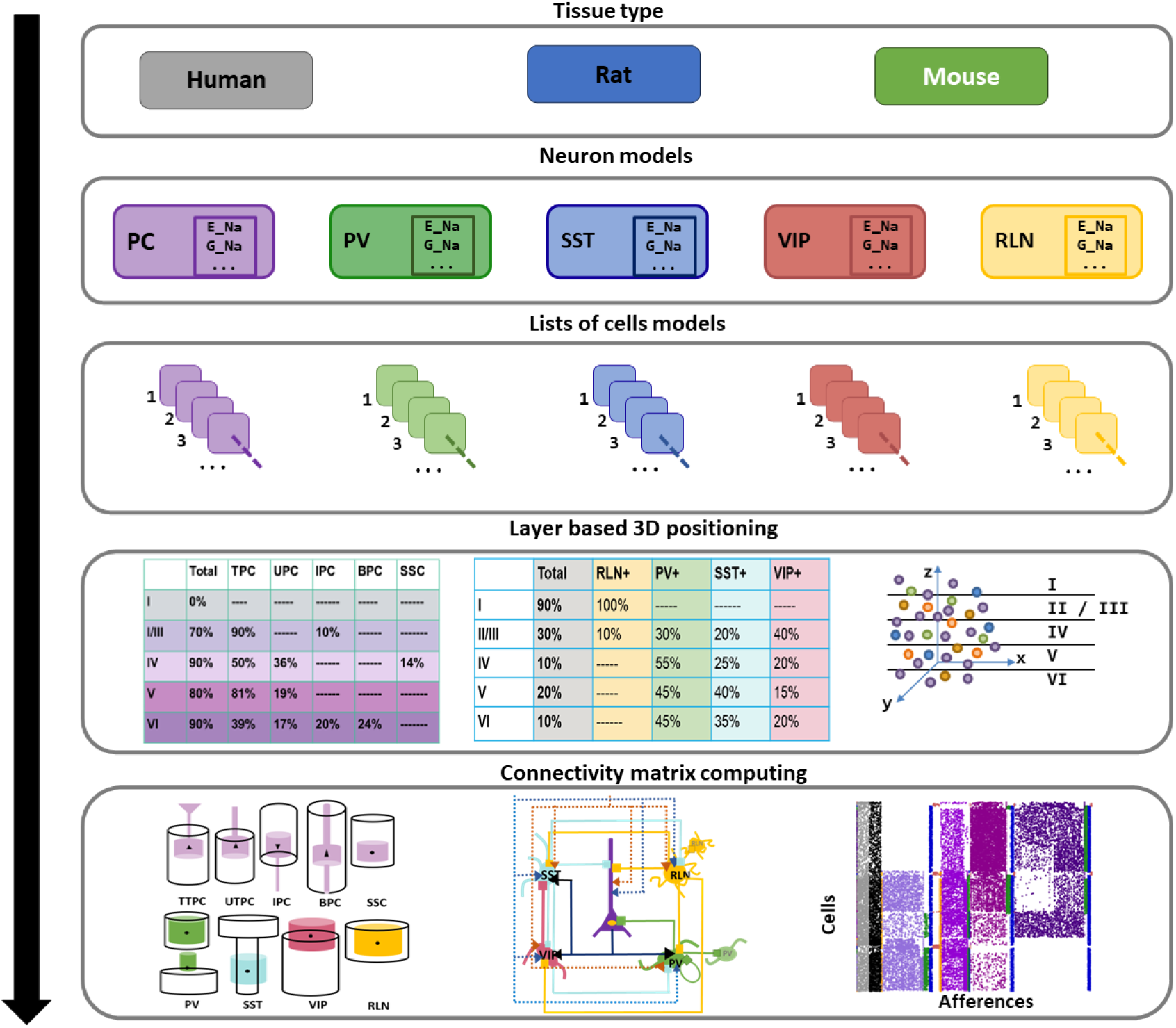
The cortical model architecture and implementation sequence. The first step is to select the tissue type from human, rat, or mouse. Then depending on the tissue type, the number, subtypes, and distribution of cells per layer are defined. For each type of neuron, the corresponding list of cells is created. Types include Principal Cells (PCs) and interneurons. The PCs are further divided into Tufted (TTPCs), Untufted (UTPCs), Inversed (IPCs), bipolar (BPCs) pyramidal cells and Spiney Stellate Cells (SSs). The interneurons Included Parvalbumin (PV), Somatostatin (SST), Vasoactive intestinal peptide (VIP), and Reelin (RLN) expressing interneurons. The cells are then placed into the predefined 3D volume of the cortical tissue respecting the corresponding distribution per layer. Finally, the synaptic connectivity matrix is computed by respecting the predefined rules of connectivity.

- Tissue type selection: three cortical types are included in NeoCoMM. These tissue types are human, rat, and mouse. For each type, the geometrical configuration of the cortical layers and cells (layer thickness, column volume, cell type density, distribution, and the morphology of cells) is modified accordingly using the default values obtained from the literature (experimental recordings).
- Neuron models: The software includes the majority of cell types present in the cortical tissue. These types are Principal Cells (PCs) and interneurons. PCs are further divided into Tufted (TTPCs), Untufted (UTPCs), Inversed (IPCs), bipolar (BPCs) pyramidal cells and Spiney Stellate Cells (SSs). Interneurons included Parvalbumin (PV), somatostatin (SST), vasoactive intestinal peptide (VIP), and Reelin (RLN) expressing interneurons.
- List creation: In this step, the main cell type lists are created. The PCs are modeled using a three-compartment simplified model ([11]) and the interneurons a one-compartment model ([8]). The default parameters of each cell type were adjusted to match experimental recordings of firing rates.
- 3D positioning: The cells’ placement is done using the best candidate algorithm [12] by respecting their distribution in each layer.
- Computing of the connectivity matrix: The synaptic connectivity matrix is computed first by computing the overlapping of neurites between source and target cells and then by respecting the predefined afference matrix that provides the distribution of afferences for each cell type. The default afference matrix is based on the literature [13, 14, 15, 16, 17, 11] and can be modified by the user.

Once the neural network architecture has been defined and created, NeoCoMM can be used to run simulations. The block diagram of a NeoCoMM simulation is portrayed in Figure 4.

**Figure 4:**
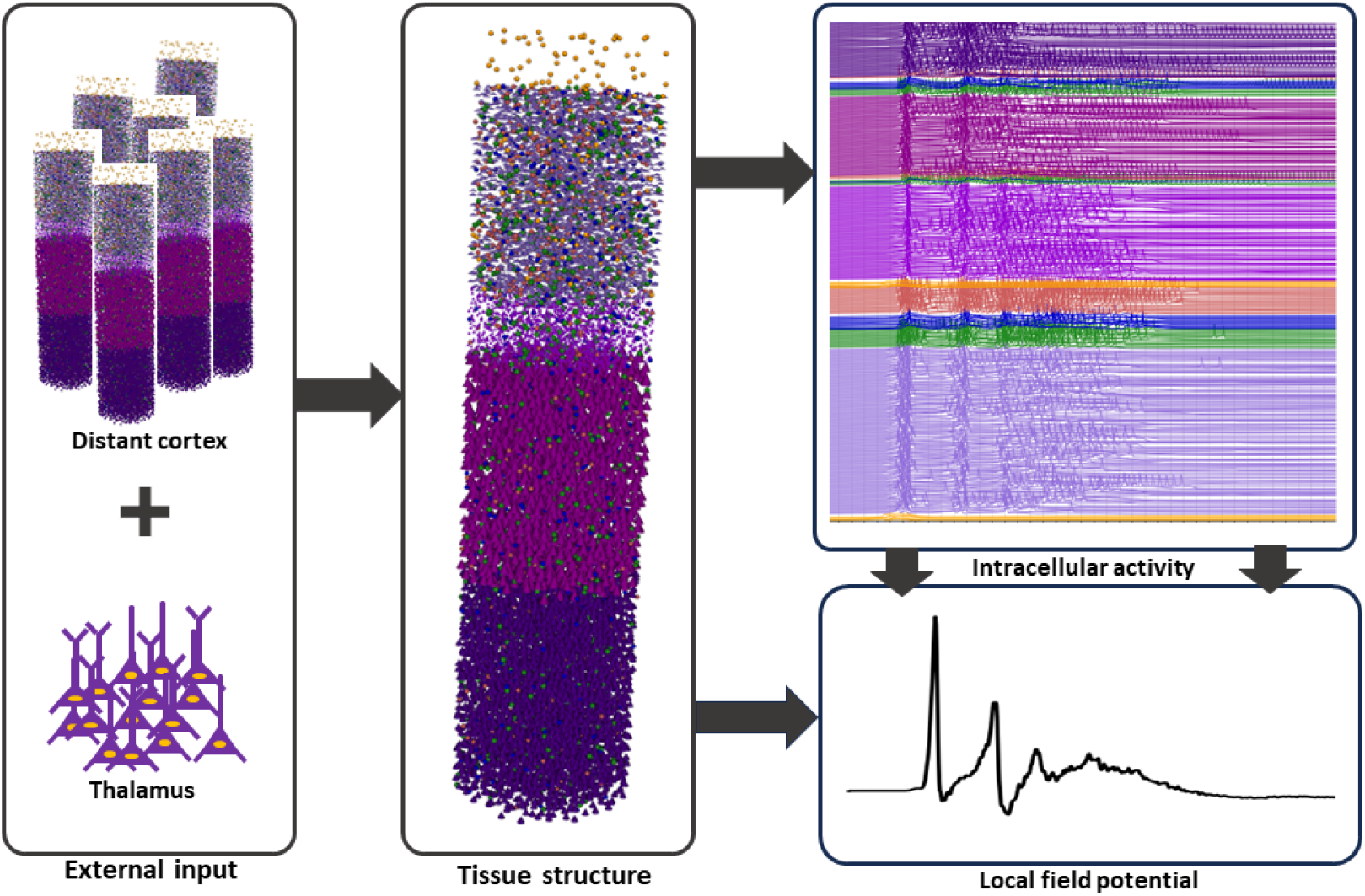
Simulation block diagram. To run a simulation, the external input must first be defined. The cortical tissue receives input from the distant cortex and the thalamus, and the parameters of each input can be adjusted by the user. In this example, we used a highly synchronized input from the distant cortex to elicit an epileptic double spike and wave from a hyperexcitable cortical column. The software will run the simulation respecting the given configuration and will provide the intracellular activity of individual cells and the local field potential recorded by the defined electrode configuration.

### 2.2. Software functionalities

Thanks to the GUI, the user can easily adapt the simulation for a specific purpose.

- Tissue creation: The dimensions and shape (cylinder, cube, or parallelepiped) of the cortical column, along with the thickness of each layer, can be adjusted.
- Selection and arrangement of neurons: The type and quantity of neurons in each layer can be adjusted, enabling the creation of diverse neuron densities that may differ based on the specific brain region. The neurons automatically adapt to the type of organism the model is attempting to simulate (human, rat, or mouse).
- Synaptic connections: A connectivity matrix can be fine-tuned, with its values either representing either a fixed number of connections between two types of neurons (which can be of the same type and layer), or a percentage of the total number of neurons involved.
- Adjusting neuron parameters: Each neuron is represented using a Hodgkin and Huxley formalism. The parameters of an individual neuron or a group of similar neurons can be adjusted through a dedicated interface featuring 3D interactions. This interface simplifies the selection and tuning of internal model variables, as well as the creation of specific clusters.
- External input: Inputs originating from the Thalamus and distant cortex are independently adjustable and can be configured to deliver a volley of action potentials either in a single burst or periodically. In addition, the noise can be seeded to replicate identical inputs, facilitating the examination of parameter effects under consistent noise conditions.
- Simulation file: The simulation can be saved to and reloaded from using a file in a text format, enabling modifications to be made to a simulation at a later time.
- Graphical representation: The GUI offers various plots for visualizing the results, including the transmembrane voltage of each neuron (at the soma) and the resulting local field potential, which is a projection of every postsynaptic potential onto an implanted electrode. The tissue is also depicted in 3D using a VTK 3D view, which is also used for selecting and modifying neuron parameters. The connectivity matrix is represented in a colorful scatter plot to convey the complexity of interconnectivity. Finally, stimulations applied to external inputs (Thalamus and distant cortex) can also be depicted.

## 3. Illustrative examples

An illustrative video of the use of NeoCoMM to simulate interictal spike and wave is provided. In this video, we show how the model can be used to simulate the realistic cellular and extracellular activity of a human neural network. We also show the different panels that can be used to test different hypotheses and to adjust the network, stimulation, and recording electrode parameters.

## 4. Impact and conclusions

The NeoCoMM software is unique in its ability to provide a compromise between complexity and realism. Compared to highly complex, time and resource-consuming microscale models, NeoCoMM represents a simplified computational approach while also incorporating physiological and biophysical realism. One of the main advantages it offers is the ability to test different hypotheses related to cellular processes and synaptic dynamics in a healthy or pathological context [5]. The user can easily install the software to explore and test different scenarios and neural network configurations. Different types of simulations can be performed using NeoCoMM. Examples of such simulations can be the study of the impact of synaptic connectivity of certain neural groups on the firing rate of neurons, the synchronization level of the external stimulation input from the distant cortex, and the thalamus effect on the local field potential characteristics, the action potential count and its relation to the different voltage-dependent channels parameters… In another area, NeoCoMM can be valuable in explaining observed patterns recorded from subjects with different brain pathologies as in [11]. In this recent study, using NeoCoMM we explained the underlying dynamics of interictal events generation. This proves that NeoCoMM can help answer research questions that cannot be resolved experimentally. NeoCoMM can also be used to test and optimize therapeutic options for brain disorders. We are currently, using NeoCoMM to study the impact of transcranial current stimulation on the recorded epileptic events (spikes, seizures, HFOs…). It allows us to explore the neuromodulatory acute effect of tCS on the individual cell’s activity in a network setting that is still unknown. The Software also permits the testing of different configurations of stimulation (intensity, type, and frequency) and their underlying effect on the pathological cell responses. Another area in which the Software can be used to improve the recording of certain events by optimizing the electrode designs using the already provided electrode-tissue interface model. This model can be adapted to test out different electrode configurations (recording surface, electrode placement, orientation, material…) and their impact on the simulated LPF such as in [18]. Finally, this Software provides a starting point for a larger framework that can simulate multiple cortical columns and be adjusted to represent larger network dynamics.

## 5. Acknowledgements

This project has received funding from the European Research Council (ERC) under the European Union’s Horizon 2020 research and innovation program (grant agreement No 855109)

